# Multi-platform plasma proteomics reveals orthogonal metabolic signatures distinguishing elite athletic phenotypes

**DOI:** 10.64898/2026.02.05.704017

**Authors:** Vincent Albrecht, Marta Murgia, Daniela Schranner, Martin Schönfelder, Johannes B Müller-Reif, Martin Halle, Henning Wackerhage, Matthias Mann

## Abstract

Exercise confers profound health benefits, yet the molecular mechanisms linking physical activity to health and longevity are incompletely known. Here we applied three mass spectrometry (MS)-based and one aptamer-based proteomics workflows to elite athletes with contrasting metabolic phenotypes, sampled before and after maximal exhaustive exercise. MS detected larger effect sizes and resolved isoforms; aptamers extended proteome coverage but with unannotated proteoform biases. Acute exercise induced coordinated platelet degranulation, neutrophil activation, and extracellular matrix turnover, with peptide topology analysis providing direct evidence for vesicular release. Chronic adaptations organized along two orthogonal axes: a muscle mass gradient marked by hypertrophy signaling and attenuated systemic inflammation, and an oxidative capacity gradient characterized by metabolic health-associated proteins (APOA4, IGFBP2, ITLN1) and dampened IGF-I signaling. Exploratory biological age analysis suggested younger adipose age in athletes. The plasma proteome provides an integrated readout of exercise adaptation, linking cardiorespiratory fitness to metabolic health and healthy aging.

## Introduction

Physical activity ranks among the most potent interventions for extending human health span. Regular endurance or resistance exercise substantially reduces cardiovascular disease, type 2 diabetes, cancer, and neurodegenerative disease risk – benefits often rival or exceed those achieved with pharmacological therapies^1–4^. Cardiorespiratory fitness, quantified as maximal oxygen uptake (VO_2_max), is among the strongest predictors of longevity: each 10 mL/kg/min increase associates with a 45-day life expectancy extension across 46 years of follow-up^5^. Similarly, muscular fitness scores such as higher grip strength are associated with reduced mortality^6^. Yet the molecular mechanisms underlying these systemic benefits remain incompletely understood.

Blood plasma provides a unique window into the integrated physiological state of an organism. As the medium through which organs communicate, plasma contains proteins secreted by virtually every tissue – myokines from contracting muscle, adipokines from fat, hepatokines from liver, or exerkines as proteins secreted during exercise – creating a circulating record of inter-organ crosstalk^7–9^. Exercise acutely and chronically remodels this plasma proteome, but investigating the scope of these changes has been constrained by technological limitations. Mass spectrometry (MS)-based approaches offer unbiased discovery and the ability to distinguish protein isoforms but face dynamic range challenges in plasma; affinity-based platforms provide broader nominal coverage but cannot resolve proteoforms and may suffer from cross-reactivity. Most studies employ only a single analytical platform, leading to a fragmented picture of exercise-induced change^10,11^. Acute responses are frequently dominated by abundant leakage proteins like creatine kinase and myoglobin with chronic adaptations inferred from targeted inflammatory panels, and little integration across temporal and phenotypic dimensions.

Elite athletes represent physiological extremes of human metabolism – the anabolic state of natural bodybuilders, the glycolytic power of sprinters, the oxidative capacity of endurance athletes – shaped by unique genetics or talent and years of discipline-specific training. Whether these divergent metabolic phenotypes manifest as distinct circulating protein signatures that integrate into coherent biological programs remains unexplored.

To broadly characterize the elite athlete plasma proteome we here applied four complementary workflows – three MS-based approaches (undepleted plasma, acid precipitation, nanoparticle corona) and an aptamer-based platform (Illumina/SOMAmer technology) – to plasma from highly anabolic, glycolytic and oxidative athletes versus recreationally active controls sampled before and after maximal exhaustive exercise. This multi-platform strategy exploits the distinct physicochemical selectivity of each method: acid precipitation implicitly enriches lower-abundance soluble species; nanoparticle coronas preferentially capture membrane-associated and vesicular proteins; aptamer arrays provide broad coverage with orthogonal physicochemical biases^12,13^. Integration across workflows addresses limitations that have constrained previous studies. As an exploratory analysis, we applied the OrganAge framework, which uses organ-enriched plasma proteins to estimate biological age for eleven organ systems^14^, to ask whether athletic phenotypes associate with biological age estimates.

Our findings reveal that acute exhaustive exercise induces rapid plasma proteome remodeling through platelet degranulation, neutrophil activation, and extracellular matrix turnover. Chronic adaptations establish distinct baseline signatures along two orthogonal axes: a muscle mass gradient characterized by ECM remodeling and attenuated inflammation, and an oxidative capacity gradient marked by metabolic health-associated proteins. This analysis suggests younger adipose biological age in athletes, with convergence on proteins identified independently by MS. The plasma proteome thus serves as an integrated readout of exercise adaptation with implications for understanding how physical activity may preserve metabolic health.

## Results

### Complementary proteomics workflows provide validated coverage of the athlete plasma proteome

We applied three complementary MS-based plasma proteomics approaches and one aptamer-based platform to samples from elite male athletes – sprinters as a model for high glycolysis (n=8), natural bodybuilders as a model for high anabolism (n=9), and endurance athletes as a model for high oxidative metabolism (n=11) – including a German champions – collected before and after a graded cycle exercise test to subjective exhaustion; and recreationally active controls (n=7) (Methods, Supplementary Data 1, Fig. 1a). The proteome depths of the MS-based workflows reflected their enrichment principles: undepleted plasma (NEAT) analysis identified a total of 1,099 protein groups; perchloric acid enrichment (PCA-N) extended coverage to 1,459 proteins by precipitating high-abundance species whereas bead-based enrichment (BEADS) achieved 2,009 protein groups through nanoparticle corona formation. The aptamer-based Illumina Protein Prep 6K platform (hereafter ‘Illumina’) provided 6,831 targets – corresponding to 30% of protein coding genes with bias toward secreted proteins and extracellular domains. Note that 41% of targets showed signal above twice the limit of detection across all samples and that certain analytes with non-linear behavior were excluded from standard reporting (Supplementary Data 1, Fig. 1b).

**Fig. 1:**
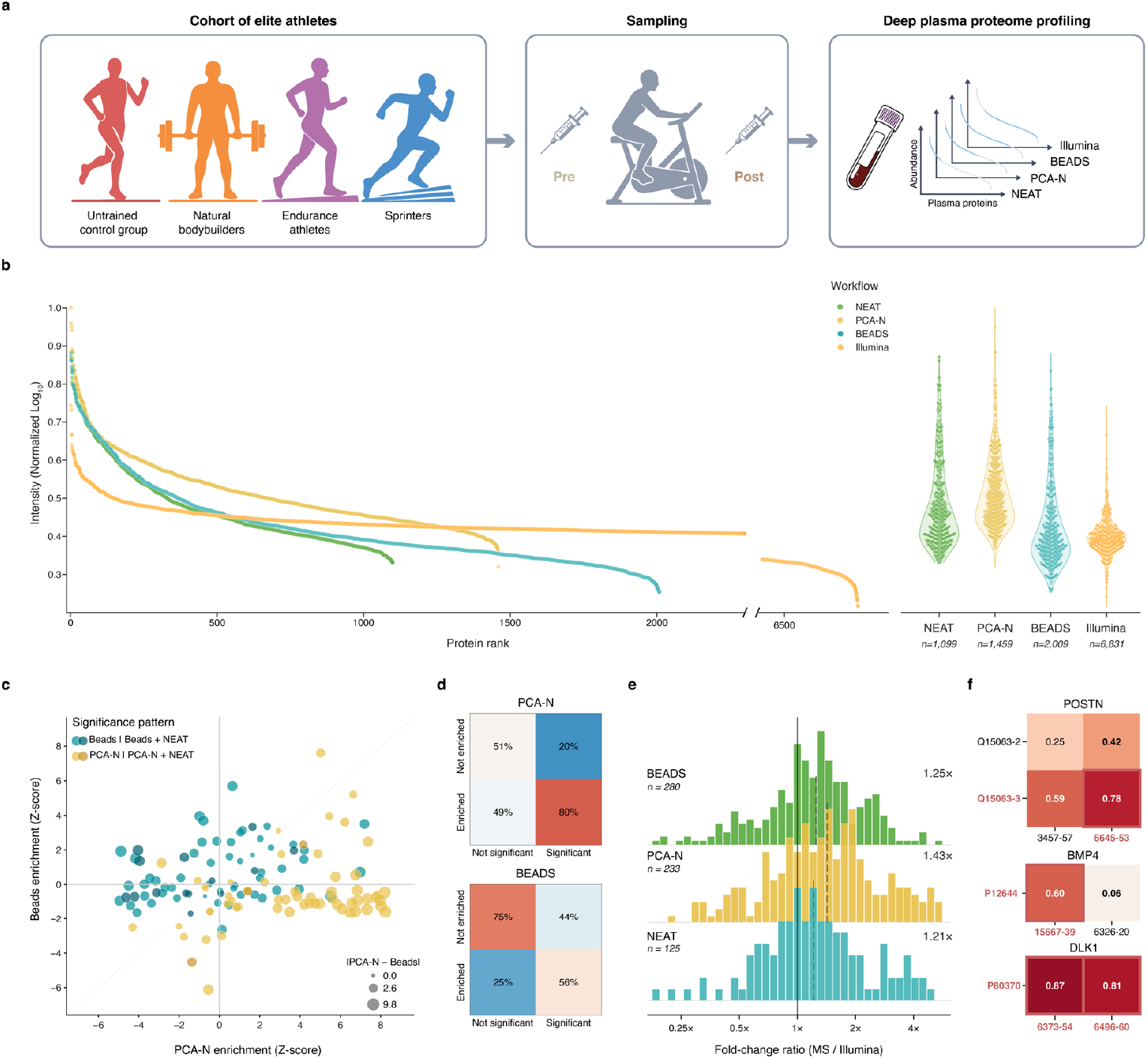
Complementary proteomics workflows provide validated coverage of the athlete plasma proteome. **a**, Study design schematic. Four athlete groups (controls, n=7; sprinters, n=8; bodybuilders, n=9; endurance athletes, n=11) before (Pre) and after (Post) exercise on a cycle ergometer. Plasma was processed using three MS-based workflows – NEAT (undepleted plasma), PCA-N (perchloric acid precipitation) and BEADS (nanoparticle-based enrichment) – and one aptamer-based platform (Illumina). **b**, Proteomic depth of different plasma proteomics workflows. Protein abundance rank curves with violin plots showing abundance distributions across workflows. **c**, Relationship between workflow-specific enrichment and statistical significance. Protein enrichment in PCA-N (x-axis) versus BEADS (y-axis) relative to NEAT plasma expressed as Z-scores. Teal: significant in BEADS or BEADS+NEAT; yellow: significant in PCA-N or PCA-N+NEAT; dot size: absolute enrichment difference between workflows (|PCA-N – BEADS|). Significance based on acute exercise and chronic training models. **d**, Enrichment-significance association. Proportion of proteins enriched (relative to NEAT) among significant versus non-significant proteins. **e**, Cross-platform fold-change comparison for genes quantified by both MS and IPP. Fold changes based on chronic training model. **f**, Spearman correlation heatmaps between MS protein isoforms and SOMAmer aptamers for multi-mapping genes. Red labels: significant; red borders: significant in both platforms. POSTN and BMP4 (BEADS); DLK1 (PCA-N).

To validate that MS workflow-specific enrichment translates to genuine biological signal we examined the relationship between protein enrichment relative to NEAT and statistical significance in the exercise response. Significant proteins in PCA-N were preferentially enriched (80% versus 49% of non-significant proteins), consistent with acid precipitation enhancing detection of biological effects otherwise masked by high-abundance species. These proteins – such as functionally important glycoproteins – exhibit lower technical variance, higher fold changes and greater statistical power. BEADS showed a more NEAT-like pattern, with non-enriched proteins rarely reaching significance (75% non-enriched among non-significant versus 44% among significant) – reflecting the equilibrium binding and surface competition inherent to corona formation (Fig. 1c,d; Supplementary Data 2).

To assess cross-platform concordance, we compared fold changes for proteins quantified by both MS and Illumina. MS consistently detected larger effect sizes: 65–69% of overlapping proteins showed greater absolute fold changes (P< 0.01, Wilcoxon test), translating directly to increased statistical power with more MS-only than Illumina-only significant proteins across all workflows (15 versus 7, 19 versus 10, and 18 versus 12 for NEAT, PCA-N, and BEADS, respectively). Proteins reaching significance on both platforms showed complete directional agreement and strong correlation (ρ up to 0.87), establishing ground truth for target validation (Fig. 1e, Extended Data Fig. 1a,b; Supplementary Data 2). From this reference, we could deduce aptamer-isoform relationships in cases of apparent platform discordance: for POSTN, significance of one isoform and one aptamer, combined with their strong correlation (ρ = 0.78 versus 0.25–0.59 for other pairings), identified the specific aptamer-proteoform match; for BMP4, divergent correlations between two aptamers and the same protein (ρ = 0.60 versus 0.06) revealed epitope-dependent recognition. Where both aptamers targeted the same proteoform, as for DLK1, all three measurements reached significance with ρ > 0.81 (Fig. 1f, Extended Data Fig. 2; Supplementary Data 2). These findings establish that aptamer reagents are often blind to proteoforms – information that MS inherently provides given sufficient peptide coverage.

Gene ontology analysis confirmed distinct subcellular origins across workflows (Fig. 1e). All four platforms enriched extracellular and secretory terms, as expected for plasma. NEAT and BEADS additionally captured lipoprotein particles, while PCA-N depleted these species but uniquely enriched signaling receptors and cell substrate junctions. BEADS and Illumina each captured additional unique terms including muscle-associated proteins. These orthogonal enrichment profiles confirm that the workflows provide complementary biological windows rather than redundant depth, enabling dissection of acute and chronic exercise responses.

### Acute exercise remodels the plasma proteome via platelet, matrix and immune activation

Samples collected before and five minutes after the cycle ergometer test revealed substantial plasma proteome remodeling across all workflows, with markedly different yields following stringent filtering (q < 0.001, |log_2_FC| ≥ 0.5): 2 significantly altered proteins in NEAT, 8 in PCA-N and 41 in BEADS (Fig. 2a; Extended Data Fig. 3a,b; Supplementary Data 3). Gene ontology enrichment revealed compartment-specific origins: BEADS showed strong enrichment for platelet alpha-granule and secretory granule lumen while PCA-N predominantly captured collagen-containing extracellular matrix proteins – consistent with the capacity of nanoparticle coronas to preserve vesicular structures that acid precipitation disrupts (Extended Data Fig. 3c).

**Fig. 2:**
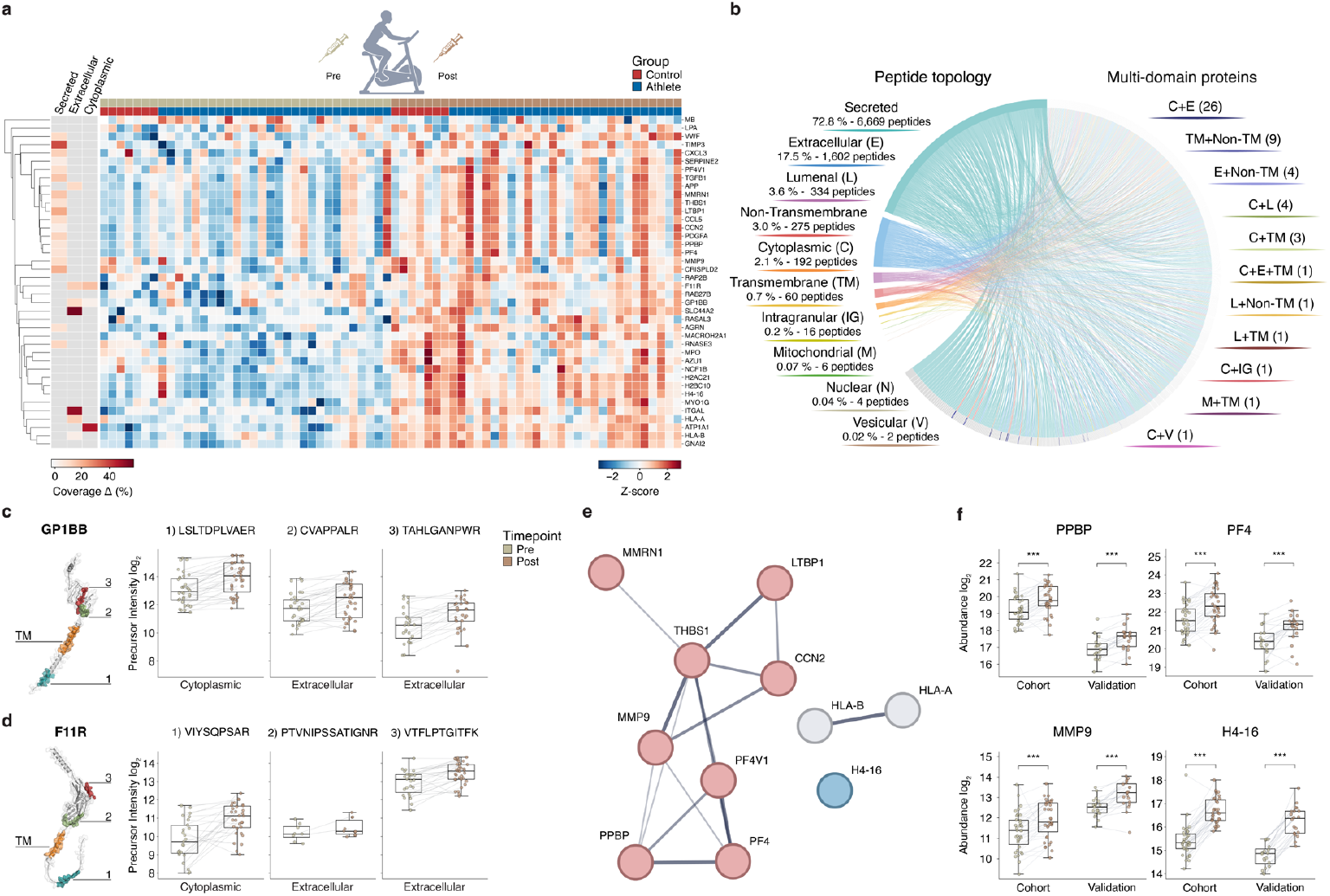
Acute exhaustive exercise rapidly remodels the plasma proteome. **a**, Heatmap of significantly changed proteins following acute exhaustive exercise in the BEADS workflow (q < 0.001, |log_2_FC| ≥ 0.5; 41 proteins). Rows represent proteins; columns represent samples ordered by timepoint (pre: beige; post: brown) and group (control: red; athlete: blue). Left annotation: peptide topology coverage by category. **b**, Peptide topology analysis. Circos plot shows subcellular origin of detected peptides based on UniProt annotations. Left: peptide distribution by compartment. Right: multi-domain proteins with peptides from multiple compartments. **c**,**d**, Dual-topology peptide detection for transmembrane proteins (**c**) GP1BB and (**d**) F11R. Left: protein structure with detected peptides mapped by domain. TM, transmembrane domain. Right: peptide abundance by topology and timepoint. **e, f**, Validation of exercise-responsive proteins across discovery and validation cohorts.(**e**) Interaction network of consistently elevated proteins across MS workflows. (**f**) Representative proteins from each functional module (BEADS data shown).

MS quantifies proteins by aggregating their identified peptides, enabling investigation of release mechanisms through topology mapping. Of 9,160 peptides with UniProt domains annotations across 1,398 proteins in BEADS, 73% originated from secreted proteins and 18% from extracellular domains with similar distributions across all three workflows (Fig. 2b; Extended Data Fig. 3d,e). Notably, dozens of proteins yielded peptides from both intracellular and extracellular compartments, indicating release of intact membrane-spanning proteins. This is exemplified by GP1BB and F11R, single-pass transmembrane proteins whose cytoplasmic and extracellular peptides showed coordinate post-exercise increases (Fig. 2c,d) – direct evidence for release within platelet-derived microparticles or extracellular vesicles^15^ rather than purely proteolytic ectodomain shedding.

Validation in an independent cohort (n=20, including female participants) confirmed eleven proteins as consistently elevated after this extremely short acute exercise – representing a core acute response signature that generalizes across cohorts and sexes (Fig. 2e,f, Extended Data Fig. 3f) clustered into three functional modules: platelet alpha-granule components (PF4, PF4V1, PPBP, MMRN1, THBS1),extracellular matrix factors (MMP9, CCN2, LTBP1) and immune signaling molecules (HLA-A, HLA-B, H4-16), as confirmed by enrichment analysis (Extended Data Fig. 3g). The aptamer platform independently recovered the platelet core of this signature (PF4, PPBP, THBS1) alongside MMP9, providing orthogonal confirmation of the dominant acute response module (Extended Data Fig. 3h).

The five platelet-derived proteins indicate that degranulation is a major contributor to the acute exercise secretome^16,17^. PF4 and PF4V1 are chemokines released upon activation^18^; PPBP is the precursor of beta-thromboglobulin, a classical *in vivo* activation marker^19^; MMRN1 binds coagulation factor V^20^; and THBS1 activates latent TGF-β while exerting anti-angiogenic effects^21–23^. This signature is consistent with exercise-induced shear stress and catecholamine-mediated platelet activation^24,25^.

The ECM triad reflects initiation of matrix turnover. MMP9 degrades type IV collagen^26^, CCN2 is a mechanosensitive matricellular protein rapidly induced by mechanical loading^27^ and LTBP1 sequesters latent TGF-β in the matrix and regulates its bioavailability^28^, suggesting engagement of TGF-β-linked remodeling programs in post-exercise repair.

Detection of MMP9, stored in granules of polymorphonuclear leukocytes, together with histone H4 suggests activation of circulating innate immune cells. Extracellular histones are released during NETosis, whereby activated neutrophils expel chromatin decorated with granule proteins. Strenuous exercise induces NET formation, with cell-free DNA originating primarily from mature neutrophils^29^.

These three modules are mechanistically interconnected. Activated platelets release THBS1, which activates latent TGF-β complexes bound to LTBP1 – a pathway known to induce CCN2^30^. Platelet-derived chemokines recruit and activate circulating immune cells, and platelets can directly trigger NET formation by neutrophils. Together, this coordinated response – balancing tissue remodeling with controlled inflammation – prepares the organism for subsequent repair and adaptation.

### Baseline plasma proteome distinguishes athlete phenotypes along orthogonal physiological axes

Beyond acute responses, we examined whether resting plasma reflects chronic training adaptations. Linear mixed-effects models comparing athletes to sedentary controls identified 13, 20, 17 and 25 differentially abundant proteins in NEAT, PCA-N, BEADS and Illumina respectively (FDR < 0.05), enriched for muscle cell differentiation, migration and growth regulation (Extended Data Fig. 4a–g; Supplementary Data 3).

To leverage the complementary protein coverage of each workflow, we integrated 303 proteins significantly associated with athlete group (35 NEAT, 48 PCA-N, 70 BEADS, 150 Illumina;FDR < 0.05; Supplementary Data 3) using MultiOmics Factor Analysis (MOFA), achieving clear phenotype separation. Bodybuilders and controls occupied distinct regions, with endurance and sprint athletes positioned intermediately (Fig. 3a; Extended Data Fig. 4h; Supplementary Data 4).

**Fig. 3:**
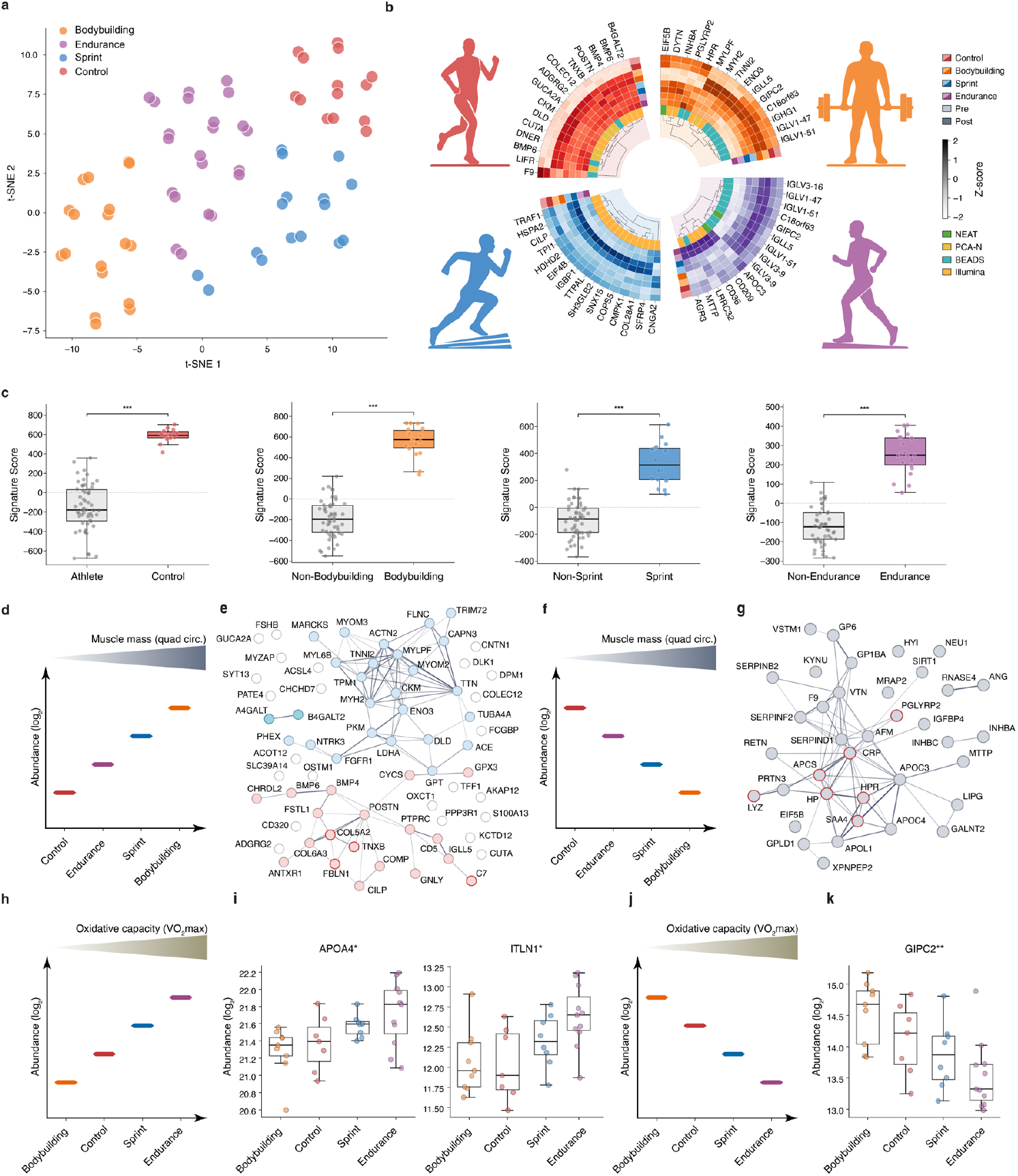
Baseline plasma proteome distinguishes athlete phenotypes along orthogonal physiological axes. **a**, t-SNE projection of integrated proteomics data (303 proteins across four workflows). **b**, Circular heatmap of top-ranked proteins per athlete group. Inner track indicates source workflow. **c**, Composite signature scores comparing each group against all others. ***P< 0.001. **d–g**, Muscle mass axis (based on quadriceps circumference). (**d**,**f**) Group ordering schematics (**e**) Network for 75 proteins increasing with muscle mass; circled node indicate liver fibrosis biomarker overlap^43^. (g) Network for 37 proteins decreasing with muscle mass; circled proteins indicate inflammatory response components. **h–k**, Oxidative capacity axis (based on VO_2_max). (**h**,**j**) Group ordering schematics. (**i**,**k**) Representative proteins with monotonic trends. *P < 0.05, **P < 0.01.

Group importance scores derived from MOFA latent factors revealed workflow-specific contributions: MS workflows dominated the bodybuilding, endurance and control signatures (11, 11 and 9 of 15 top-ranked, respectively), with BEADS and PCA-N contributing most substantially; the sprint signature derived exclusively from Illumina (Fig. 3b; Extended Data Fig. 4i). The MS approach resolved distinct isoforms – FGFR1 splice variants (P11362-16 and P11362-21) contributed independently to the control signature, and PGLYRP2 proteoforms (M0R2W8 and Q96PD5) ranked separately among bodybuilding-associated proteins – information inaccessible to affinity platforms (Extended Data Fig. 5a–d). Composite signature scores significantly distinguished each group from all others, with bodybuilding and control showing the clearest separation (Fig. 3c).

These phenotype differences organized along two orthogonal physiological axes: a muscle mass gradient (control < endurance < sprint < bodybuilder, based on quadriceps circumference) and an oxidative capacity gradient (bodybuilder < control < sprint < endurance, based on VO_2_max).

### Circulating proteins scale with muscle mass and resistance training

We next examined the protein signatures defining each axis. Linear mixed-effects modelling identified 75 proteins increasing progressively along the muscle mass axis (control < endurance < sprint < bodybuilder; Fig. 3 d,e; Supplementary Data 3), with functional enrichment for sarcomere organization and muscle structure development (Extended Data Fig. 4j).

The most striking finding was coordinate elevation of fast-twitch sarcomeric components – MYLPF, TNNI2, MYH2, ACTN2, MYOM2 andMYOM3; glycolytic enzymes PKM, ENO3 and LDHA; and the muscle-specific creatine kinase CKM^31,32^. This profile is consistent with an increasing fraction of fast muscle fibers from endurance athletes to bodybuilders, combined with larger relative volume of fast compared to slow fibers. The appearance of these intracellular proteins in plasma likely reflects both increased fast muscle mass and exercise-induced sarcolemmal permeability^33^ – positioning the plasma proteome as a non-invasive window into muscle quantity and fiber type composition. TRIM72 (MG53), which nucleates sarcolemmal membrane repair by translocating to injury sites and facilitating vesicle fusion, showed progressive elevation^34^; suggesting that this gradient reflects adaptive membrane repair capacity in response to chronic mechanical loading.

Proteins involved in muscle stem cell regulation and myogenesis followed the same pattern: DLK1, critical for postnatal muscle hypertrophy^35^; BMP4 and BMP6, positive regulators of muscle mass signaling through Smad1/5/8^36,37^; FGFR1, a regulator of satellite cell proliferation^38,39^; and MARCKS, whose calpain cleavage is required for myoblast fusion^40^ – collectively, candidate biomarkers for satellite cell activation and fusion during hypertrophy and repair. ACE, whose D allele associates with greater hypertrophic responses to resistance training, also increased along the gradient^41,42^.

A prominent ECM subset – FBLN1, CHRDL2, TNXB, COL5A2, COMP, POSTN, CILP and C7– mediates matrix assembly, collagen fibril organization and modulation of TGF-β signaling. Intriguingly, several of these proteins have been independently identified as plasma biomarkers of liver fibrosis progression, with FBLN1 showing the strongest correlation with fibrosis stage^43^. This convergence suggests that these proteins report ECM remodeling activity rather than tissue-specific processes, emphasizing the need for context-dependent biological interpretation. In athletes, elevated ECM proteins likely reflect myotendinous junction remodeling and mechanical load adaptation, whereas in liver disease they mark fibrosis. The anti-fibrotic properties of CILP in TGF-β–driven contexts^44^ may contribute to maintaining adaptive rather than pathological ECM responses.

### Attenuated inflammation and enhanced metabolic efficiency characterize trained athletes

The 37 proteins that decreased along the muscle mass axis were enriched for acute phase response, lipid metabolism and hemostasis (Fig. 3f,g; Extended Data Fig. 4k; Supplementary Data 3). Innate immune components showed coordinate suppression, including acute-phase reactants used as biomarkers of systemic inflammation. CRP showed the expected inverse relationship with lean mass, consistent with anti-inflammatory effects of chronic exercise^45,46^. This pattern also characterized serum amyloid P component (APCS)^47^, serum amyloid A4 (SAA4)^48^, haptoglobin (HP)^49^ and its related protein (HPR)^50^, lysozyme (LYZ)^51^ and peptidoglycan recognition protein 2 (PGLYRP2)^52^. Proteinase 3 (PRTN3), a neutrophil serine protease whose plasma levels correlate with cardiovascular risk factors, and VSTM1/SIRL-1, an inhibitory myeloid receptor regulating neutrophil oxidative burst, similarly decreased. Athletes with greater muscle mass thus exhibit reduced inflammatory burden – a phenotype associated with lower cardiometabolic risk that may represent an underappreciated benefit of muscle mass-increasing resistance training.

Proteins elevated in insulin-resistant states followed the same pattern: APOC3, APOC4, APOL1, MTTP, AFM, RETN and LIPG – each elevated in type 2 diabetes and metabolic syndrome^53–56^ – declined progressively from controls to bodybuilders, consistent with the dominant role of skeletal muscle in glucose disposal^57^. The liver-derived phospholipase GPLD1, a novel prediabetes biomarker whose plasma levels rise alongside fasting hyperglycemia^58^, similarly decreased, as did GALNT2, possibly reflecting its role in adipocyte maturation^59^. Kynureninase (KYNU), an enzyme of the inflammation-activated tryptophan catabolic pathway, also decreased – consistent with enhanced kynurenine clearance by exercise-trained muscle^60^.

Six coagulation and fibrinolysis proteins (F9, SERPIND1, SERPINF2, SERPINB2/PAI-2,GP1BA, VTN)^61–63^ decreased in parallel, together with glycoprotein VI (GP6), the major platelet collagen receptor implicated in arterial thrombosis, suggesting reduced thrombotic potential and enhanced fibrinolytic capacity – cardiovascular protective adaptations consistent with lower disease risk in physically active populations^64^.

Strikingly, key negative regulators of muscle mass converged on this axis. IGFBP4, which sequesters IGF-I and limits its bioavailability^65,66^, showed pronounced inverse association with muscle mass, as did activin A (INHBA) and inhibin βC (INHBC)—TGF-β superfamily members that signal through ActRIIB to phosphorylate SMAD2/3 and function synergistically with myostatin as negative regulators of muscle mass. Pharmacological activin blockade reverses wasting in cachexia models and ligand traps have recently received clinical approval^67,68^. The plasma proteome thus captures the principal relays governing muscle mass: diminished inhibitory TGF-β/activin signaling combined with enhanced IGF-I bioavailability.

### Oxidative capacity associates with a longevity-like plasma signature

Orthogonal to the muscle mass axis, 35 proteins scaled monotonically with estimated oxidative capacity (bodybuilder < control < sprint < endurance; Fig. 3h–k; Supplementary Data 3). Eighteen increased and 17 decreased with VO_2_max, converging on metabolic flexibility and vascular adaptation.

Three proteins positively associated with oxidative capacity – APOA4, IGFBP2 and ITLN1 – have each been independently linked with human longevity. APOA4 alleles are enriched in centenarians^69–72^; ITLN1 (omentin-1) parallels adiponectin, elevated in centenarians and predictive of reduced mortality^73^; and IGFBP2 limits IGF-I bioavailability^74–76^, mirroring the dampened IGF-I signaling linked to lifespan extension across species^77,76^. This attenuation of IGF-I signaling was reinforced by decreased GIPC2, a PDZ-domain adaptor that facilitates IGF-I receptor signaling^78^ – indicating reduced pathway activity at both ligand (IGFBP2) and receptor-proximal (GIPC2) levels, consistent with endocrine profiles associated with extended lifespan in model organisms and centenarian studies^77,79^.

APOA4 promotes fatty acid oxidation while enhancing glucose disposal through dual hepatic and adipose mechanisms^80^, and its increase alongside ITLN1 delineates a metabolic flexibility signature consistent with the ‘athlete’s paradox’ of preserved insulin sensitivity despite elevated intramuscular lipid stores^81^. This profile distinguishes the endurance phenotype from the glycolytic/anabolic phenotype of bodybuilders.

Vascular adaptation markers also scaled with oxidative capacity, consistent with higher capillary density of slow-oxidative compared to fast-glycolytic muscle fibers. ESAM, an endothelial junction molecule that promotes capillary sprouting, and PLXNB2, the functional receptor for angiogenin, both increased – reflecting VEGF-driven angiogenesis during endurance training.

Eight immunoglobulin components and the acute-phase protein ITIH4, Fc-binding glycoprotein (FCGBP), and granulysin (GNLY) decreased with increasing oxidative capacity, consistent with lower basal inflammatory tone and exercise-associated immunomodulation^82^.

### Organ-specific biological age reveals distinct effects of acute exercise and chronic training

To integrate protein-level findings into a systems-level framework, we applied the OrganAge models^14^ using Illumina (Fig. 4a; Supplementary Data 5). This approach estimates biological age for 11 organ systems using plasma proteins with organ-enriched expression. Organ-specific age gaps derived from these models have been associated with future disease risk and mortality in independent population-based cohorts^14,83,84^.

**Fig. 4:**
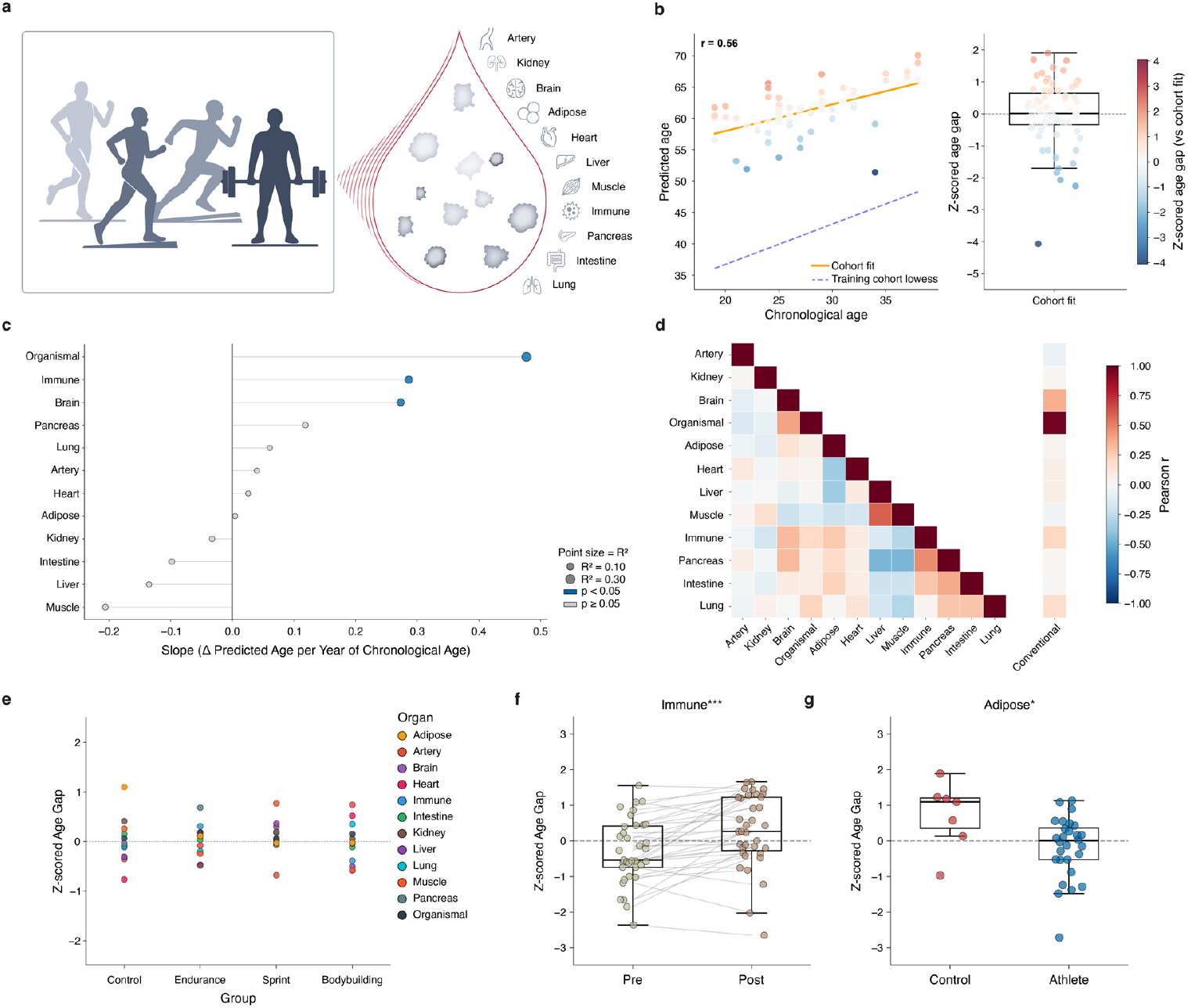
Organ-specific biological age reveals distinct effects of acute exercise and chronic training. **a**, Schematic of OrganAge framework for 11 organ systems plus organismal and conventional models. **b**, Predicted versus chronological age for the Organismal model (r = 0.56). Solid line: cohort-specific fit; dashed line: training cohort LOWESS curve. **c**, Organ model calibration; slope of predicted versus chronological age per organ, point size indicates R^2^. **d**, Inter-organ correlation heatmap (mean r = 0.037). **e**, Organ age profiles by training modality. Dashed line indicates population mean. **f**, Acute exercise effect on immune biological age: Δ = +0.58 SD, Cohen’s d = 0.77, ***P < 0.001. **g**, Chronic training effect on adipose biological age. Δ = −0.92 SD, Cohen’s d = −1.05, *P = 0.043.

The original models were trained on elderly cohorts (∼75 years). Given our young cohort (19– 38 years), we recalibrated age gaps using within-cohort regression, yielding z-scored residuals where positive values indicate biologically “older” and negative values “younger” organ age relative to same-aged peers. Despite the narrow age range, predicted organ ages correlated with chronological age (r = 0.56 for the Organismal model; Fig. 4b,c), and low inter-organ correlations (mean r = 0.037; Fig. 4d) confirmed that organs age heterogeneously within individuals – a pattern extending to young, fit populations, with distinct profiles across training types (Fig. 4e).

Acute exercise most strongly affected immune biological age (Δ = +0.58 SD, p < 0.001, Cohen’s d = 0.77; Fig. 4f), consistent with transient innate immunity activation^85,86^ and aligning with the post-exercise platelet and neutrophil activation observed in our proteomic data (Fig. 2).

Chronic training effects were most pronounced for adipose tissue: athletes exhibited significantly younger biological age than controls (Δ = −0.92 SD, p = 0.043, Cohen’s d = −1.05; Fig. 4g). The Adipose OrganAge model comprises only five adipose-enriched proteins, among them ITLN1 (omentin-1) – which MS independently identified as increasing along the oxidative capacity axis (Fig. 3). This convergence of protein-level and organ-level findings implicates adipose tissue as a key locus of training-associated biological age differences.

## Discussion

This study establishes the plasma proteome as an integrated readout of both acute physiological responses to exercise and the chronic metabolic phenotypes that distinguish athletes as a consequence of genetic predisposition and discipline-specific training.

A multi-workflow proteomic strategy was essential to resolve these effects: acid precipitation enriched muscle-derived intracellular proteins while depleting vesicular structures; nanoparticle coronas preferentially captured platelet-derived microparticles and membrane-associated species; aptamer arrays provided orthogonal coverage enabling organ-level aging analysis. Direct cross-platform comparison revealed that MS consistently captures larger effect sizes through superior dynamic range, while aptamer reagents exhibit implicit proteoform selectivity – with concordance depending critically on isoform-aptamer matching. That such selectivity remains unannotated provides a path toward enhanced aptamer characterization; that MS resolves proteoforms inherently, combined with ongoing advances in depth, positions mass spectrometry to ultimately deliver both sensitivity and comprehensive coverage. Matching the proteoform complexity of the human proteome accessible to MS will require aptamer panels far exceeding current gene-level coverage. No single platform would have revealed the full scope of exercise-induced proteome remodeling – underscoring that workflow selection fundamentally shapes biological conclusions in plasma proteomics.

The acute response to a bout of graded cycle exercise to exhaustion reflects coordinated activation of interconnected systems: platelet degranulation, neutrophil mobilization and extracellular matrix turnover. Peptide topology analysis provided direct molecular evidence for vesicular release, with transmembrane proteins yielding peptides from both cytoplasmic and extracellular domains – a signature inconsistent with proteolytic shedding alone^87^. That this inflammatory mobilization manifests as a transient elevation in immune biological age suggests that biological aging clocks may, in part, capture reversible immune activation states rather than exclusively reflecting irreversible senescent processes. This interpretation contrasts with the chronic, unresolved inflammation characteristic of inflammaging^88^ and aligns with exercise as a hormetic stimulus: an acute physiological stressor that, through repeated cycles of activation and efficient resolution, promotes systemic adaptive benefits^89^.

Chronic training adaptations organized along two orthogonal axes. The muscle mass gradient captured hypertrophy signaling (DLK1, BMP4/6, ACE), ECM remodeling and attenuated systemic inflammation – collectively establishing a milieu that favors anabolic adaptation while reducing cardiometabolic risk. The coordinated reduction in coagulation factors and enhanced fibrinolytic capacity provides molecular grounding for the well-documented inverse relationship between physical activity and cardiovascular disease^64^. That several ECM proteins elevated in resistance-trained athletes overlap with liver fibrosis biomarkers^43^ illustrates a central challenge in circulating biomarker interpretation: these proteins report generalized matrix remodeling, with biological significance determined by clinical context rather than the molecules themselves.

The oxidative capacity axis yielded one of the study’s most striking findings: convergence on molecular programs associated with human longevity. APOA4, IGFBP2 and ITLN1 – each independently linked to exceptional longevity or reduced mortality^69–72^ – were elevated in endurance athletes, alongside coordinated attenuation of IGF-I signaling at both ligand and receptor levels. This signature recapitulates the hormonal profile associated with lifespan extension across species^73,77^. The organ-level analysis reinforces this interpretation: athletes exhibited younger adipose biological age, with ITLN1 – a visceral adipose-derived adipokine that enhances insulin-stimulated glucose uptake and exerts anti-inflammatory effects ^90–92^ – identified independently by MS as increasing along the oxidative capacity axis. Circulating ITLN1 levels are reduced in obesity and insulin resistance but increase following aerobic training, even in absence of weight loss^93,94^. This convergence of protein-level and organ-level findings suggests that regular training preserves adipose tissue in a metabolically favorable state, providing a plausible link between cardiorespiratory fitness and the slower pace of biological aging associated with reduced mortality^95^.

These findings reframe how we interpret plasma proteomics. Circulating biomarkers have traditionally served to detect pathology – elevated CRP signals inflammation, cardiac troponins indicate myocardial damage. Our data demonstrate that the plasma proteome equally identifies systems functioning exceptionally well: longevity-associated signatures, younger organ biological age and attenuated inflammatory profiles represent molecular evidence of physiological optimization. Plasma proteomics thus provides a comprehensive readout of human physiological status across the spectrum from pathology to peak performance – a perspective with implications for clinical diagnostics and human performance optimization alike.

These findings highlight the importance of studying exercise and non-pathological phenotypes such as those of athletes to establish the context specificity of circulating protein biomarkers. This study directly tests whether proteins widely interpreted as pathological biomarkers are also influenced by athletic training or acute exercise. Creatine kinase illustrates this challenge: elevated levels indicate tissue injury in myocardial infarction but also rise transiently after exercise due to skeletal muscle leakage without pathological significance. Distinguishing disease-related biomarkers from those reflecting adaptive physiological stress is essential for accurate interpretation of plasma proteomic signals.

Several limitations warrant acknowledgment. Our cohort comprised young males, limiting generalizability; sample sizes, particularly for controls (n=7), constrain statistical power for subgroup analyses; and the cross-sectional design cannot distinguish training-induced adaptations from genetic predisposition toward both athletic phenotype and favorable proteomic profile. The immunoglobulin changes associated with oxidative capacity, while consistent with training-induced immunomodulation, could reflect confounding factors in our modest sample. The single post-exercise timepoint captures immediate responses but not recovery dynamics, and the OrganAge framework, while validated in large cohorts^14,83,84^, represents an extension beyond its original elderly training population. A key outstanding question is the relative contribution of genetics versus training: longitudinal studies tracking proteomic trajectories through training initiation, detraining, or modality switching – combined with twin studies discordant for athletic training – would help resolve how much of the athlete proteome reflects acquired adaptation versus innate predisposition, with implications for exercise prescription in clinical populations.

The plasma proteome provides accessible, minimally invasive readouts of systemic adaptation state that could inform personalized training prescription and monitor response to exercise interventions. The identification of longevity-associated signatures offers mechanistic insight into the well-established relationship between cardiorespiratory fitness and lifespan^4^. Whether these proteomic patterns predict long-term health outcomes in athletic and general populations remains to be determined, but the convergence of protein-level and organ-level findings on pathways implicated in healthy aging positions the plasma proteome as a promising tool for understanding – and potentially optimizing – the health benefits of physical activity.

## Methods

### Study cohorts

#### MetaExtreme cohort

The MetaExtreme cohort comprised healthy, trained male athletes representing three distinct training modalities – endurance athletes (n=11), natural bodybuilders (n=9) and sprinters (n=8) – alongside untrained controls (n=7). Recruitment built upon an earlier study ^96^ and followed identical inclusion criteria and pre-sampling protocols. All participants met stringent inclusion criteria and followed standardized pre-sampling protocols: adherence to a controlled diet on the day before testing, abstention from exercise for 24 hours, and discontinuation of dietary supplements for 48 hours prior to sampling. The study was approved by the Technical University of Munich ethics committee (#356/17S); all participants provided written informed consent.

#### MetaPerform validation cohort

The validation cohort comprised 20 healthy, endurance-trained athletes (9 women, 11 men), aged 22–42 years. Inclusion criteria required a normal body mass index (18.5–25.0 kg/m^2^), a minimum of 8 hours per week of endurance training and a VO_2_max exceeding 50 ml/kg/min for women and 60 ml/kg/min for men. Female participants were tested during the early follicular phase (days 1– 10) to minimize hormonal variation. Pre-sampling protocols matched the MetaExtreme cohort. The study was approved by the Technical University of Munich ethics committee (#203/20 S-EB); all participants provided written informed consent.

### Plasma proteomics

#### MS-based workflows

Three complementary mass spectrometry-based workflows were applied to exploit distinct physicochemical selectivity. The NEAT workflow processed undepleted plasma (1 μL) by standard reduction, alkylation and trypsin/LysC digestion^12^. The PCA-N workflow used perchloric acid precipitation (5 μL plasma) followed by neutralization to enrich low-abundance species while depleting albumin and immunoglobulins^12^. The bead-based workflow employed functionalized nanoparticles (OmniProt, Westlake Omics) to form protein coronas that preferentially capture membrane-associated and vesicular proteins^13^. All workflows were semi-automated using the Bravo liquid handling platform (Agilent). Digested peptides (200 ng) were loaded on Evotips following the manufacturer’s protocol and separated on an Evosep One system (60 samples-per-day method, 21 min gradient) using an 8 cm Aurora Rapid column (150 μm ID; IonOpticks) coupled to an Orbitrap Astral mass spectrometer (Thermo Fisher Scientific) operating in data-independent acquisition (DIA) mode. The instrument was interfaced with a FAIMS Pro device (compensation voltage −40 V). MS1 scans (380– 980 m/z) were acquired at 240,000 resolution; MS/MS scans used 3 Th isolation windows and a maximum injection time of 7 ms with 25% normalized HCD collision energy.

### Aptamer-based proteomics

Neat plasma samples (55 μL) were processed using the Illumina Protein Prep 6K workflow on an Illumina Protein Prep Automation System (Illumina Solutions Centre, Milan). Plasma was diluted 1:5, 1:200, and 1:20,000 to capture proteins across the dynamic range and incubated with SOMAmer reagent beads (28°C, 850 r.p.m., 3.5 h). Following incubation, beads were washed, captured proteins biotinylated, and protein– SOMAmer complexes photocleaved. Complexes were recaptured on streptavidin beads in the presence of a polyanionic competitor to reduce non-specific binding. Eluted SOMAmers were hybridized with sequencing probes (37°C, overnight) and used as templates for indexed library generation. Libraries were pooled and sequenced on a NovaSeq 6000; read counts were normalized and analysed using the DRAGEN Protein Quantification pipeline (v1.8.33). Data files were parsed using Canopy.

### Data processing and statistical analysis

#### MS-based proteomics data processing

Raw data were processed using DIA-NN v1.8.1 with match-between-runs enabled and an *in silico* predicted library searching against the human TrEMBL and SwissProt FASTA database (UniProt, November 2023, taxonomy ID 9606). Enzyme specificity was set to Trypsin/P with one missed cleavage allowed. Carbamidomethylation of cysteine was set as fixed modification; methionine oxidation as variable modification. Mass accuracy and MS1 accuracy were 10 ppm with scan window radius of 6. False discovery rate was controlled at 1% at the peptide-to-spectrum match level. Protein inference was performed at gene level using single-pass neural network classification. Quantification employed “Robust LC (high accuracy)” with RT-dependent cross-run normalization. Protein quantification used MaxLFQ intensities. Filtering for ≥20% valid values across samples yielded 1,099 (NEAT), 1,459 (PCA-N) and 2,009 (BEADS) proteins. For all subsequent analyses, proteins quantified in ≥70% of samples were retained. Batch effects were corrected using ComBat with temporary imputation; original missing values were restored post-correction. Intensities were log_2_-transformed prior to analysis.

#### Peptide topology annotation

Peptides were mapped to UniProt topology annotations (transmembrane domains, topological domains) based on sequence position. Coverage changes between pre- and post-exercise timepoints were calculated as the mean difference in peptide detection frequency across subjects for each domain.

#### Functional enrichment analysis

Gene set enrichment was performed using g:Profiler with Gene Ontology Cellular Components (excluding electronic annotations) and Reactome as reference databases. Significance was assessed using the Benjamini-Hochberg procedure (FDR <0.05). Results were visualized using EnrichmentMap in Cytoscape. Protein interaction networks were generated using STRING.

#### Linear mixed-effects modelling

To account for the repeated-measures structure (pre- and post-exercise samples from each individual), we employed linear mixed-effects (LME) models with subject as a random intercept. Models were fitted using restricted maximum likelihood (REML) estimation with the statsmodels package (Python). Two model specifications addressed distinct biological questions: (1) Acute exercise effects: Protein ∼ Timepoint + (1|Subject), testing systematic changes from pre-to post-exercise across all participants regardless of training status. (2) Chronic training adaptations: Protein ∼ Group + (1|Subject), testing for differences between groups reflecting long-term training adaptations. This model was applied to two grouping schemes: (a) athletes (bodybuilders, endurance athletes, and sprinters pooled) versus sedentary controls, and (b) the four training modalities separately (bodybuilding, endurance, sprint, control). For all models, p-values were corrected using the Benjamini-Hochberg procedure (FDR < 0.05).

#### Multi-omics factor analysis

To leverage complementary protein coverage across workflows, significantly group-associated proteins were integrated using Multi-Omics Factor Analysis (MOFA). Group importance scores were derived from latent factor loadings, and composite signature scores were computed by aggregating group importance-weighted protein abundances.

#### Monotonic trend analysis

Proteins with monotonic abundance changes along physiological axes were identified using Spearman’s rank correlation between ordered group assignments and protein abundance. Pre- and post-exercise measurements were averaged per subject prior to analysis. For the muscle mass axis, groups were ordered as control < endurance < sprint < bodybuilder. For the oxidative capacity axis, groups were ordered as bodybuilder < control < sprint < endurance (based on measured VO_2_max). Monotonic trends were considered significant at FDR < 0.05 (Benjamini-Hochberg correction) with |ρ| ≥ 0.4. Proteins were classified as “increasing” or “decreasing” based on the sign of the correlation coefficient.

### Organ-specific biological age estimation

#### OrganAge model application

Organ-specific biological ages were calculated for all 70 samples (35 subjects × 2 timepoints) using the OrganAge framework^14^ and associated Python package (https://github.com/hamiltonoh/organage), which requires the complete set of model proteins. Age estimates were generated for adipose tissue, artery, brain, heart, immune tissue, intestine, kidney, liver, lung, muscle, and pancreas, plus an organismal model using organ-nonspecific proteins.

#### Cohort-specific age gap calculation

Age gaps were recalibrated for our young cohort following the package’s recommended approach for cohort effects. For each organ model, predicted age was regressed against chronological age using ordinary least squares, and residuals were z-score transformed to yield cohort-adjusted age gaps. Positive values indicate biologically older and negative values indicate biologically younger organ age relative to same-aged peers within the cohort.

#### Statistical analysis of organ ages

Acute exercise effect (Pre vs. Post): Linear mixed-effects models with timepoint as fixed effect and subject as random intercept: AgeGap ∼ Timepoint + (1|Subject). Effect sizes were calculated as Cohen’s d for paired samples. Chronic training effect (Athletes vs. Controls): Pre and Post values were averaged per subject to obtain single independent observations, avoiding pseudo-replication. Welch’s t-test was applied given unequal sample sizes (28 athletes vs. 7 controls). Effect sizes were calculated as Cohen’s d with pooled standard deviation. Statistical significance was assessed at α = 0.05. Given that 12 organ models were tested for each question, formal multiple testing correction was not applied.

#### Software

Analyses were performed in Python 3.10 using statsmodels v0.14 for linear mixed-effects models, scipy v1.11 for statistical tests, OrganAge v1.0, and MOFA2 for multi-omics integration. Network visualization used Cytoscape v3.9 with EnrichmentMap.

## Acknowledgement

We thank our colleagues in the Department of Proteomics and Signal Transduction, Max Planck Institute of Biochemistry, for their input and support. We thank Lucia Crippa (Illumina) and Gabrielle Wheway (Illumina) for coordination and processing of the Illumina Protein Prep assay and Simon Freedman (Illumina) for support with data analysis. This study has been supported by the Max Planck Society for Advancement of Science (M.Mann). H.W. and is member of FOR 5795 which is funded by the Deutsche Forschungsgemeinschaft (DFG, German Research Foundation) (536691227). D.S. was supported by a doctoral scholarship from the German Academic Scholarship Foundation (Studienstiftung des Deutschen Volkes).

## Author Contributions

V.A. and M. Murgia conceptualized the study with input from M. Mann. V.A. performed sample preparation and mass spectrometry measurements. V.A. and M. Murgia performed data analysis and finalized study design. M. Murgia performed preliminary MS measurements and data analysis. D.S. and M.S. established the athlete cohort; M.S. coordinated sample collection; M. Halle performed physical examinations; D.S. and M.S. performed exercise physiology assessments. M. Halle, H.W., and J.B.M.-R. provided input on data interpretation.M. Mann supervised the study. V.A. generated all figures and wrote the original manuscript draft. H.W., M.H., M. Murgia, and M. Mann edited and revised the manuscript. All authors read and approved the final manuscript.

## Competing interests

M.M. is an indirect investor in Evosep Biosystems. All other authors declare no competing interests.

## Potential conflict of interest

M. M. is an indirect shareholder of Evosep. All other authors declare no competing interests.

## Supplementary Figures

**Extended Data Fig. 1:**
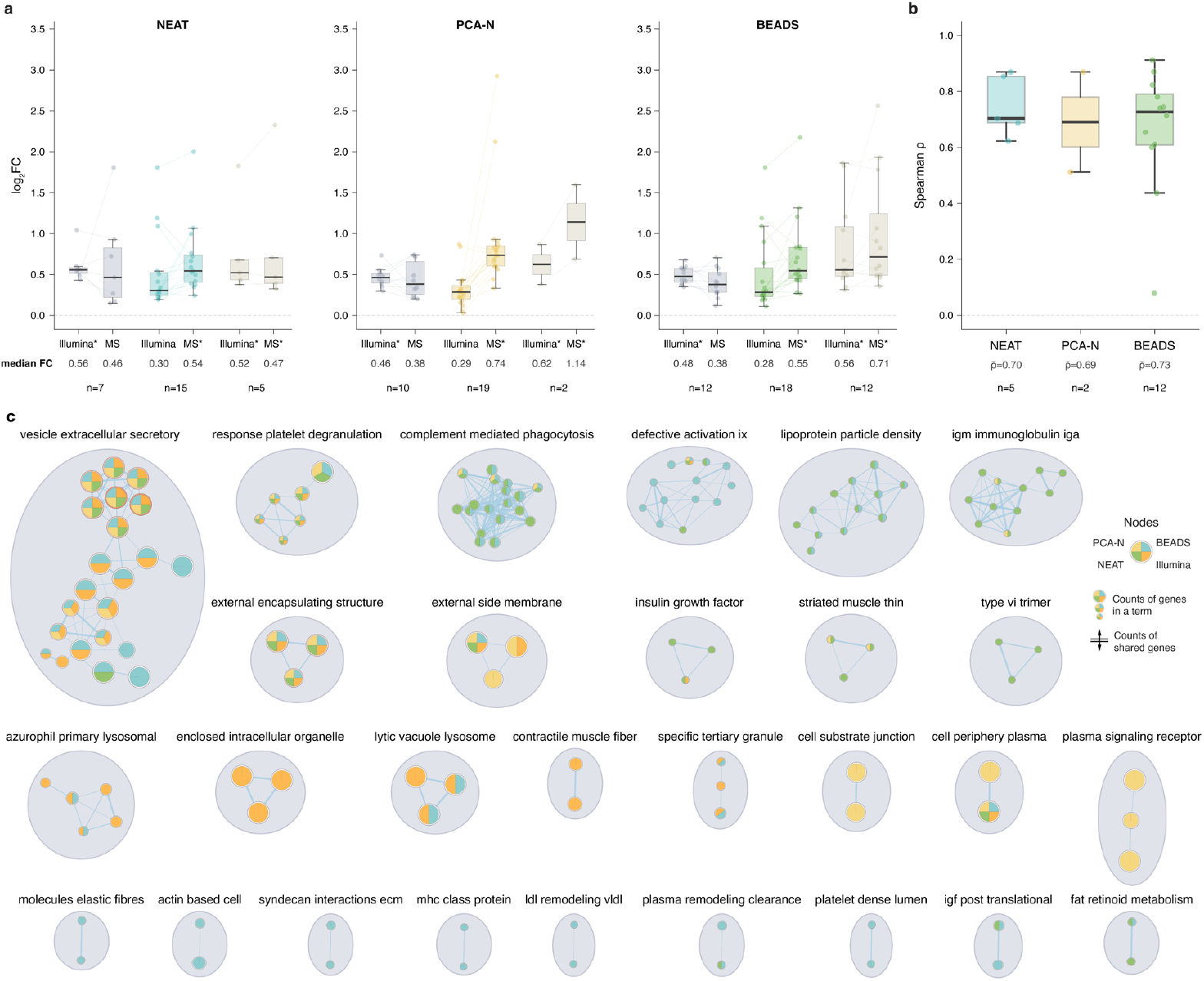
Cross-platform comparison and functional enrichment of exercise-responsive proteins. **a**, Paired fold-change comparison stratified by significance category. Asterisks indicate platform with significant proteins: Illumina only, MS only, or both. **b**, Spearman correlation of protein intensities between Illumina and MS workflows for genes significant in both platforms. **c**, Functional enrichment network of exercise-responsive proteins. Gene Ontology Cellular Component enrichment visualized using EnrichmentMap. Node size indicates gene count; color indicates significance.

**Extended Data Fig. 2:**
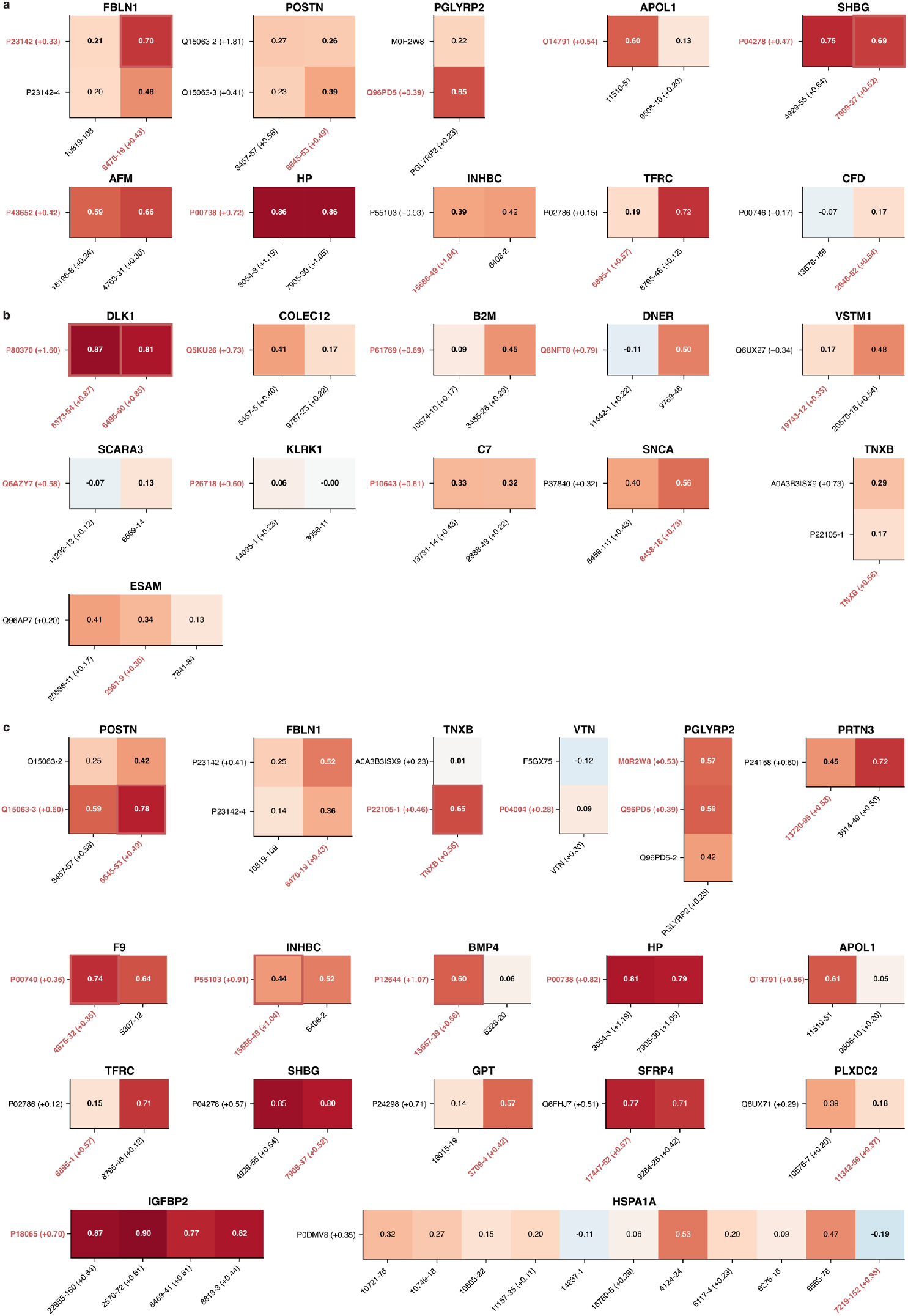
Correlation analysis of multi-mapping genes across platforms. Spearman correlation heatmaps between MS protein isoforms and SOMAmer aptamers for genes with multiple measurements. (**a**) NEAT, (**b**) PCA-N, (**c**) BEADS. Red labels indicate significant proteins (MS) or aptamers (Illumina; red borders highlight isoform-aptamer pairs significant in both platforms.

**Extended Data Fig. 3:**
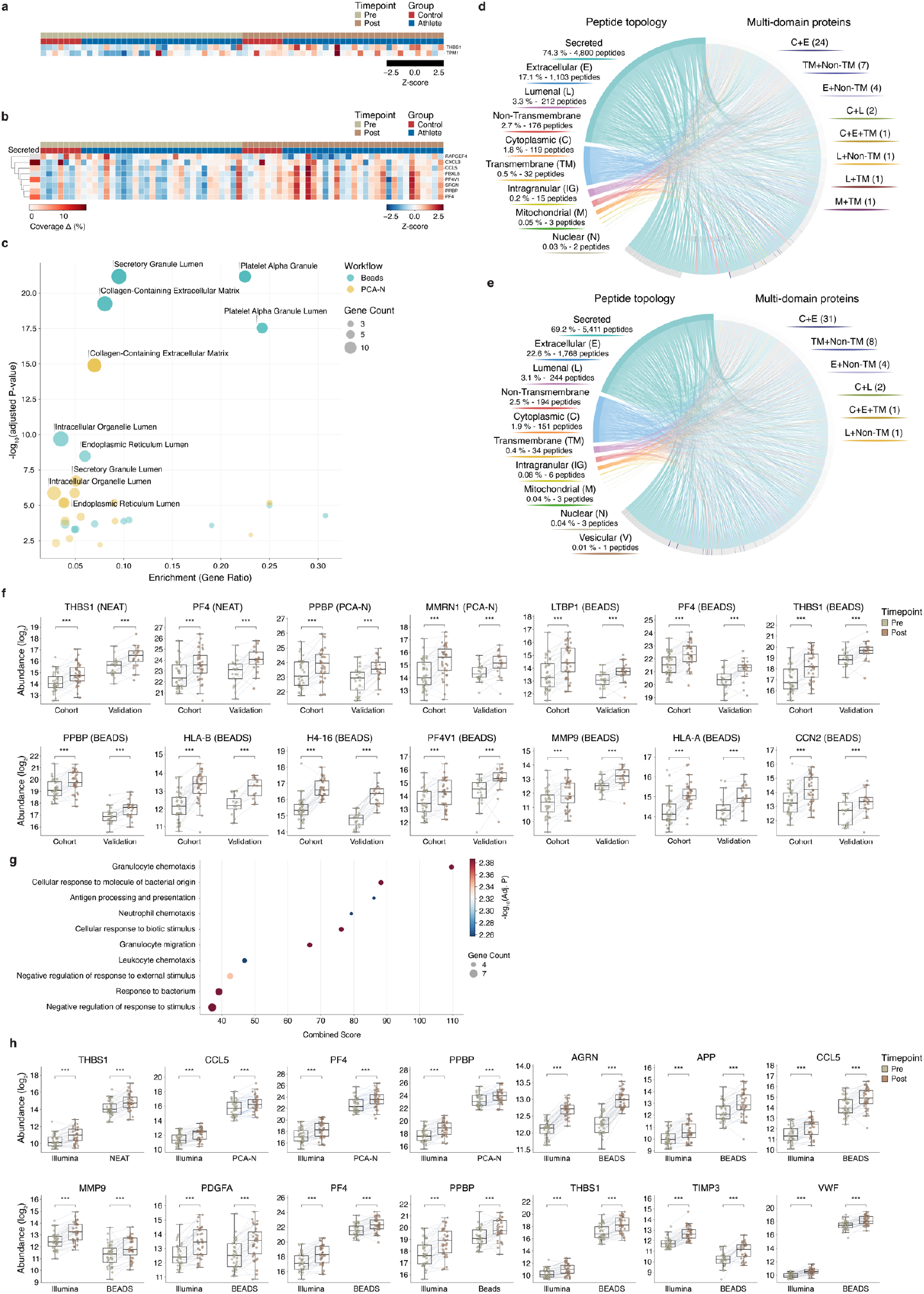
Workflow-specific acute exercise responses and validation. **a**,**b**, Heatmaps of significantly changed proteins following acute exhaustive exercise in (**a**)NEAT and (**b**) PCA-N workflows. **c**, Gene Ontology Cellular Component enrichment of significantly changed proteins by workflow. **d**,**e**, Peptide topology analysis for (**d**) NEAT and (**e**) PCA-N workflows. Left: peptide distribution by compartment. Right: multi-domain proteins with peptides from multiple compartments. **f**, Validation of exercise-responsive proteins across discovery and validation cohorts for NEAT, PCA-N and BEADS workflows. ***P < 0.001. **g**, Functional enrichment of eleven consistently elevated proteins. **h**, Cross-platform validation of acute exercise-responsive proteins in the discovery cohort. Paired comparison of protein abundance (pre versus post) for Illumina and MS workflows (NEAT, PCA-N, BEADS). ***P < 0.001.

**Extended Data Fig. 4:**
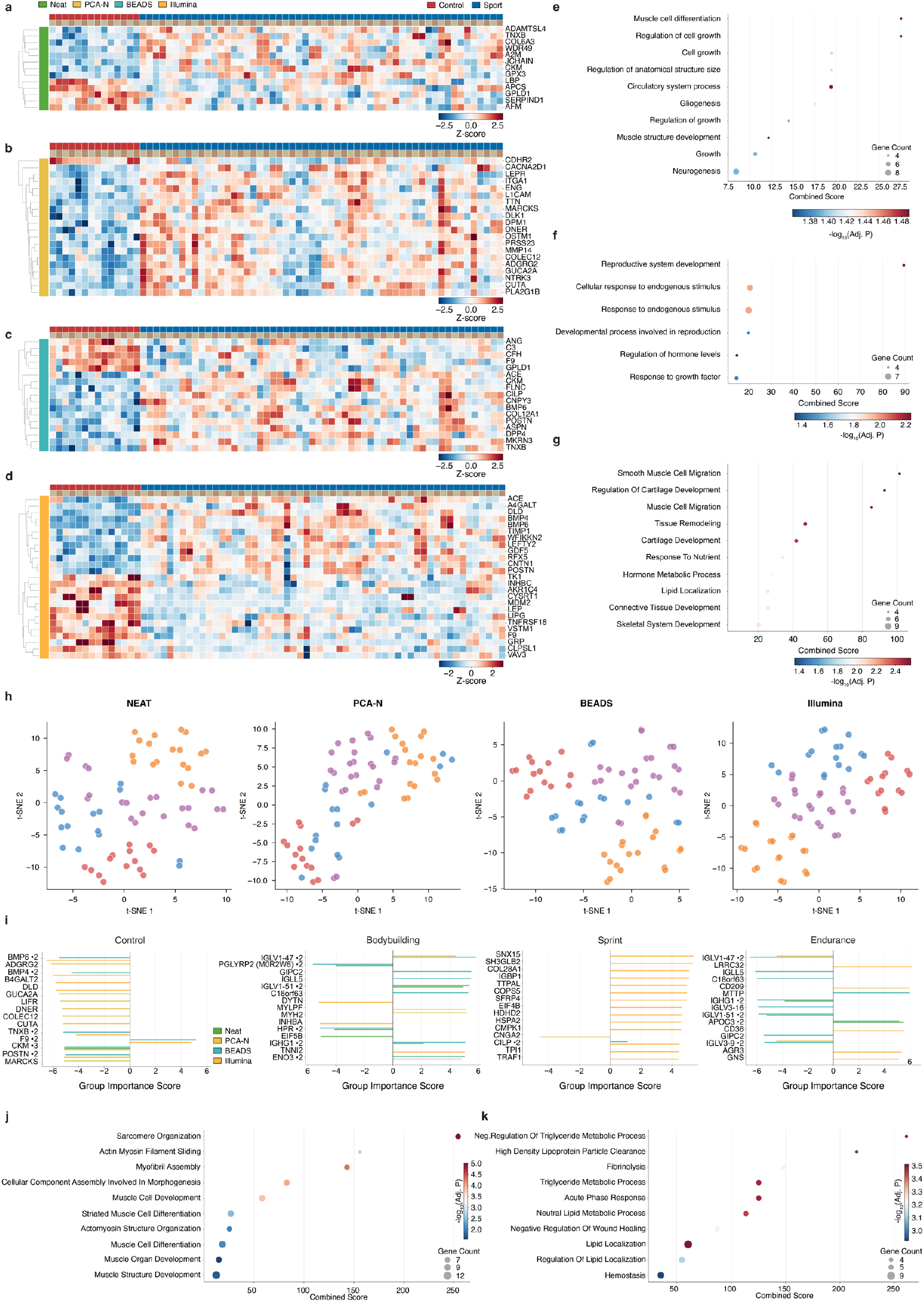
Athlete phenotype separation by workflow and integrated analysis. **a–d**, Differentially abundant proteins between athletes and controls for (**a**) NEAT, (**b**) PCA-N, (**c**) BEADS and (**d**) Illumina. **e-g**, Functional enrichment of proteins distinguishing athletes from controls for (**e**) PCA-N, (**f**) BEADS and (**g**) Illumina. **h**, t-SNE projections using workflow-specific proteins NEAT (35 proteins), PCA-N (48 proteins), BEADS (70 proteins) and Illumina (150 proteins). **i**, Top 15 proteins by group importance score for control, bodybuilding, sprint and endurance signatures. Bar color indicates source workflow; proteins detected in multiple workflows indicated with “2”. **j**,**k**, Functional enrichment of proteins along the muscle mass axis: (**j**) increasing and (**k**) decreasing with muscle mass.

**Extended Data Fig. 5:**
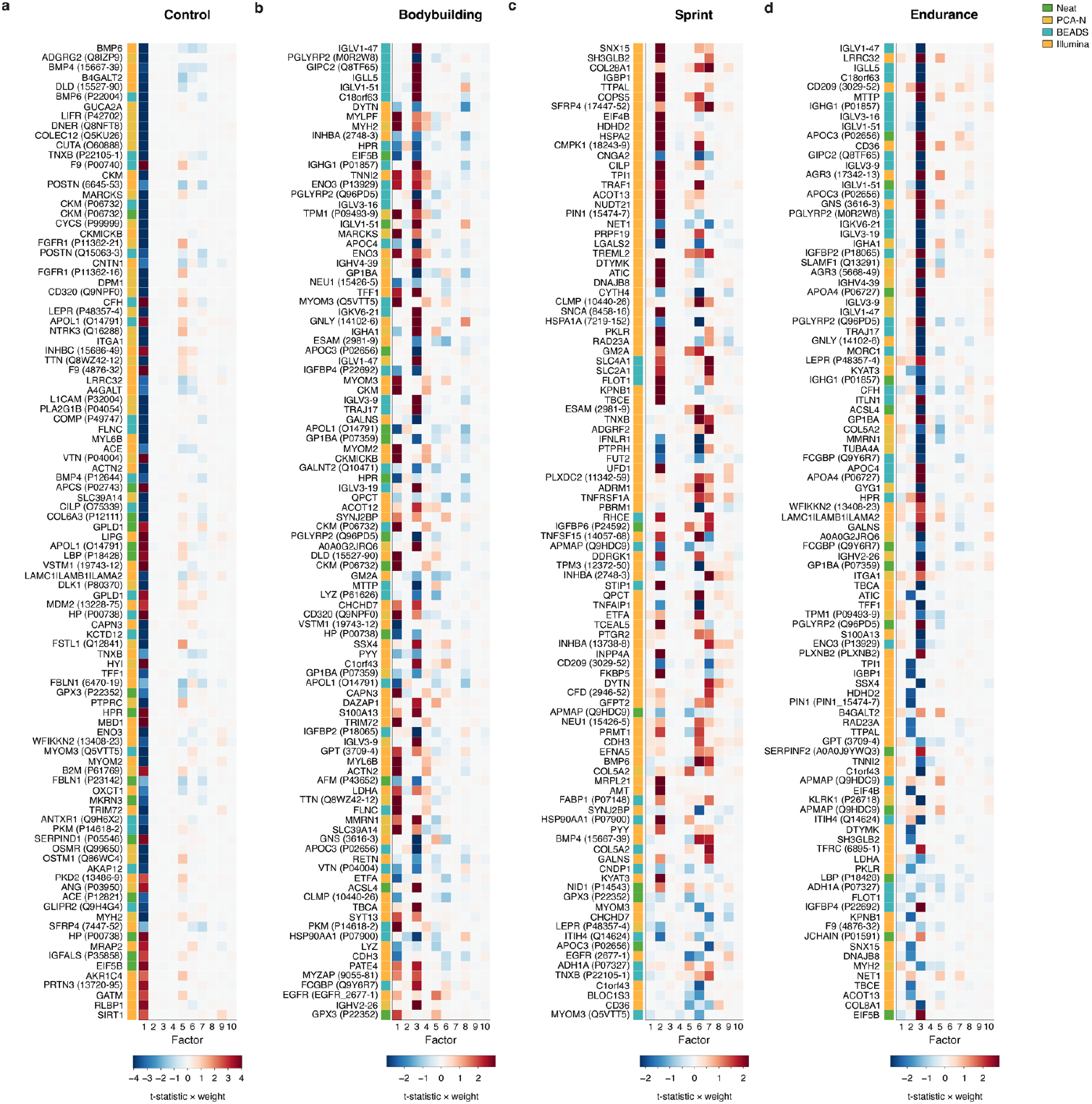
MOFA factor contribution to group-specific protein signatures. **a–d**, Protein contributions to group separation for (**a**) bodybuilding, (**b**) control, (**c**) endurance, and (**d**) sprint. Rows: proteins significant in at least one workflow (153 proteins; FDR < 0.05); columns: latent factors. Color indicates factor–group t-statistic × protein loading. Left track indicates source workflow. Proteins ordered by descending absolute group importance.

